# Structured Joint Decomposition (SJD) identifies shared dynamics across collections of biologically related multi-omics data matrices

**DOI:** 10.1101/2022.11.07.515489

**Authors:** Huan Chen, Guangyan Li, Jinrui Liu, Shreyash Sonthalia, Genevieve Stein-O’Brien, Luo Xiao, Brian Caffo, Carlo Colantuoni

## Abstract

It is necessary to develop exploratory tools to learn from the unprecedented volume of high-dimensional multi-omic data currently being produced across the field of biomedicine. We have developed an R package, Structured Joint Decomposition (SJD), which identifies components of variation that are shared across multiple matrices. The approach focuses specifically on variation across the samples/cells within each dataset while incorporating biologist-defined hierarchical structure among input experiments that can span *in vivo* and *in vitro* systems, multi-omic data modalities, and species.

SJD enables the definition of molecular variation that is conserved across systems, those that are shared within subsets of studies, and elements unique to individual matrices. We have included functions to simplify the construction and visualization of highly complex *in silico* experiments involving many diverse multi-omic matrices from multiple species. Here we apply SJD to decompose four RNA-seq experiments focused on neurogenesis in the neocortex. The public datasets used in this analysis are at [NeMO Analytics](https://nemoanalytics.org/) and can be explored at the [individual gene level](https://nemoanalytics.org/p?l=ChenEtAlSJD2022&g=DCX) or using the [conserved transcriptomic dynamics in mammalian neurogenesis](https://nemoanalytics.org/p?p=p&algo=pca&c=3a20dd81&l=ChenEtAlSJD2022) that we define here.

The SJD R package and tutorial can be found at https://chuansite.github.io/SJD.

*Contact*: [carlocolantuoni.org]

## Introduction

High-dimensional multi-omic data derived from *in vivo* tissues and *in vitro* cellular systems are now being produced at high quality and unprecedented pace. The advent of single-cell sequencing technologies has elevated archival public data resources to levels which demand creative analytical approaches to harness their greater combined discovery power. Many computational methods have been developed for the integrative analysis of high-throughput transcriptomic and epigenetic data and have been central in enabling the functional biological interpretation of complex multi-omic data. Many of these tools aim to co-register independent datasets in a unified dimension reduced space and/or rigorously define cell types that span multiple sequencing studies, often employing mutual nearest neighbor, canonical correlation analysis or related approaches (Welch et. al. 2019, Korsunsky et. al. 2019, Stuart et. al. 2019, Crow et. al. 2019).

With the goal of exploring independent studies in a more general framework, we previously developed transfer learning methods which use dimension reduction in a reference dataset to map relationships in new data via projection (projectR: Sharma et. al. 2020, Stein-O’Brien et. al. 2021). This allows variation within the new experiment to be mapped onto relative positions along dimensions discovered in the reference study, without requiring the geometric alignment of multiple datasets onto a single manifold (e.g. as is carried out in Seurat via CCA- or rPCA-based integration, Stuart et. al. 2019). Previously, we applied these projection approaches to define parallels between *in vivo* development and *in vitro* cell differentiation systems and to identify conserved transcriptomic dynamics in development across species (Kim et al 2021, Micali et al 2020). Here, we extend this flexible exploratory approach with the Structured Joint Decomposition (SJD) package to incorporate information from many datasets in the definition of dimensions of variation that are shared across multiple cellular systems under investigation.

### Approach

SJD enables the interrogation of a family of related multi-omics data matrices with common row annotation (e.g. gene, isoform, genome coordinate, etc.) to define 1] components of variation which are shared across all the datasets, 2] components common to specific subsets of the datasets, and 3] those unique to individual matrices. The resulting embeddings/scores for samples/cells in each dataset do not map to a single shared manifold, but rather to relative positions along shared dimensions. In this manner, our approach focuses specifically on within experiment variation and protects against warping of a single jointly learned manifold by between experiment variation that is often related to technological and/or batch effects.

Previously, we developed Two-stage Link Component Analysis (2sLCA; Chen et al 2022) for the decomposition of multiple matrices. We have built the rigorous statistical methodology of our 2sLCA algorithm into the SJD package, using the twoStageLCA() and twoStageiLCA() functions. In addition, we have included joint implementations of conventional principal component analysis (jointPCA()), independent component analysis (jointICA()) and non-negative matrix factorization (jointNMF()) within this same structured decomposition framework (see Supplemental Material for mathematical details of these methods and PMID: 38464021 for extensive use of the jointNMF() function). We refer to these as “joint” methods to distinguish them from the two-stage approaches. All these algorithms can be called within the SJD package and take the same arguments: 1] a list of biologically related matrices to be decomposed, 2] the requested groupings of matrices in which to search for components of variation, i.e. the grouping structure of the datasets listed in hierarchical order (e.g. all matrices, combinations of matrices that are of interest, and finally individual matrices if elements unique to individual datasets are desired), and 3] the number of components requested from each matrix grouping. 2sLCA estimates components in the matrix grouping order specified by the user, typically starting with components common to all the matrices being explored. To define components in additional matrix groupings requested by the user, 2sLCA sequentially searches spaces orthogonal to previous groupings’ components. This approach is designed to achieve the greatest separation of components across the matrix groupings requested by the user.

SJD can process matrices from any data modality that uses systematic row names that map across matrices (e.g. gene identifiers or genome coordinates). Prior to running the SJD decomposition functions, the sjdWrap() function can be used to automatically find shared rows across all the input matrices (i.e. identical row names), so rows of input matrices need not be in the same order. Because all SJD decompositions require an identical row structure for all input matrices, sjdWrap() also automatically links orthologous gene identifiers across matrices from different species using the biomaRt package which accesses the ensemble database (Durinck et al 2009). Independent of other functions in SJD, the getMatch() function called by sjdWrap() can be used to map gene identifiers in one species to orthologues in another for any application. The output of the sjdWrap() function is then used as input to any of the SJD decomposition functions. The output of SJD decompositions consists of 1] the feature loadings (i.e. row/gene weights) for as many matrix groupings as were defined by the user (the number of columns in each of these elements will depend on the number of components requested for each grouping), and 2] the sample scores (i.e. column/cell embeddings) in each dataset for each matrix grouping.

### Implementation

We illustrate the ability of SJD to define shared molecular dynamics across related matrices in a collection of four RNA-seq datasets. These datasets span *in vivo* tissues and *in vitro* stem cell differentiation, single-cell and bulk sequencing technologies, and two species to provide diverse perspectives on how neural stem cells give rise to postmitotic neurons during neurogenesis in the mammalian neocortex: 1] scRNA-seq in human fetal brain tissue (Polioudakis et al 2019), 2] scRNA-seq in mouse fetal brain tissue (Telley et al 2019), 3 and 4] bulk RNA-seq of neural induction and differentiation of human pluripotent stem cells (hPSCs; Burke et al 2020 and Ziller et al 2014). In all cases we have used the column normalized, log transformed gene level count data as processed by the original authors (see supplement for additional notes on data normalization in SJD). A vignette that guides users through this analysis and visualization can be found at https://chuansite.github.io/SJD.

Using the twoStageLCA() function, we performed a joint decomposition that defined components of variation that are shared across all four datasets, followed by components that are shared only by the two *in vitro* datasets. The first linked component (LC) shared across all four datasets is depicted in Figure 1A and reflects the primary cellular trajectory of neurogenesis across time in each of the individual datasets, showing high levels in neural stem cells (NSC) and decreasing as neurons are generated. Genes in the top 25 ranked loadings of this shared LC include proliferating NSC genes SOX2, VIM, GMNN, TOP2A and MKI67, while the bottom 25 include neuronal genes STMN2, MAPT, DCX, NRXN1 and MEF2C (Full gene loadings are in Supplemental Table 1). Gene set enrichment analysis (GSEA) of the gene loadings for this shared LC1 clearly showed this to be a trajectory leading from NSCs to post-mitotic neurons, both ends of which intersect directly with the genetics of neurodevelopmental and neuropsychiatric disease (Figure 2). Figure 1B shows the first LC common to only the two *in vitro* stem cell systems (identified in space orthogonal to the globally shared LCs). This *in vitro* specific component precisely identifies pluripotent cells that are present in the stem cell systems, but which are absent from the *in vivo* tissues at the time sampled. Genes in the bottom 25 ranked loadings of this LC (i.e. high in hPSCs) include pluripotency genes NANOG and DNMT3B (Full gene loadings are in Supplemental Table 2). Because there are no pluripotent stem cells in the *in vivo* tissues at the times sampled, many key pluripotency genes had been omitted from the *in vivo* scRNA-seq data (due to low or no expression, including OCT4/POU5F1). These genes were therefore also omitted from the joint decomposition.

**Figure 1:**
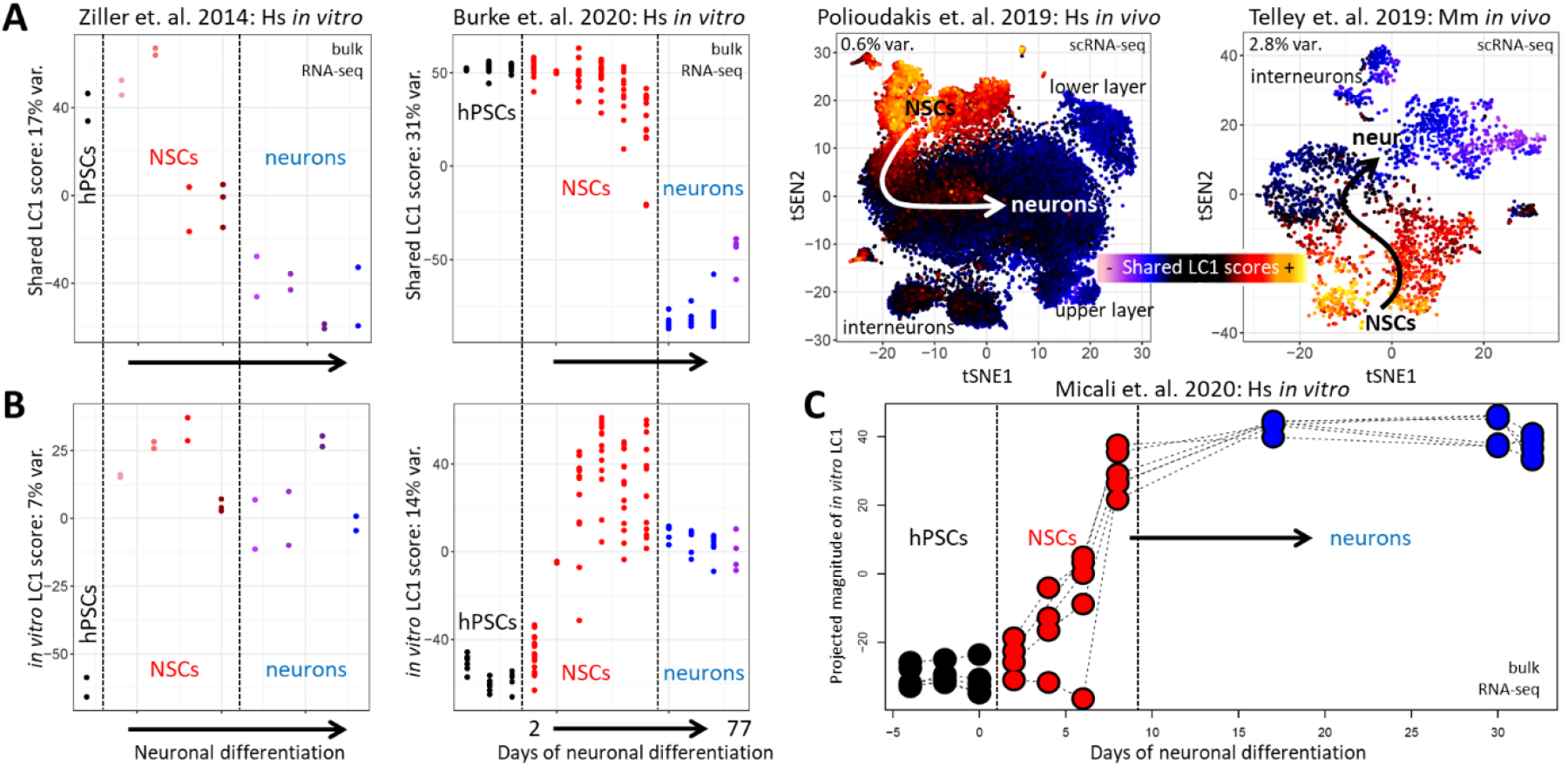
The twoStageLCA() function in SJD defines linked components (LCs) identifying conserved transcriptomic change across mammalian cortical neurogenesis. **(A)** Shared LC1 defines the primary neurogenic trajectory shared across all four datasets. In the bulk datasets LC1 scores are shown on the Y-axis (2 panels at left). In the visualization of all bulk *in vitro* datasets, the first vertical line indicates neural induction moving hPSCs toward NSCs and the second indicates the withdrawal of FGF2 (mitogen) initiating neuronal differentiation to post-mitotic neurons (in panels A, B and C). In the two scRNA-seq datasets LC1 scores are reflected in the color of individual cells plotted in the originally published tSNE coordinates (2 panels at right). In all for datasets, the descent of shared LC1 values follows the transition of neural stem cells (NSC) to neurons of the neocortex, indicating that LC1 has identified a conserved transcriptomic program in mammalian neurogenesis. **(B)** The LC1 found to be common specifically to the two *in vitro* models, identifies the human pluripotent stem cells (hPSC) that are known to be present in these stem cell systems, but absent from the *in vivo* tissues. With the exception of additional legend and axis labeling, figure panels A and B are the product of the plotting functions within SJD: SJDScorePlotter() and assemble.byComponent(). **(C)** Projection of the *in vitro* LC1 into additional hPSC differentiation data again identifies hPSCs from neural differentiation. Arrows in all panels indicate the general progression of neurogenesis. The public datasets used in this analysis are at NeMO Analytics and can be explored at the individual gene level or using the conserved transcriptomic dynamics in mammalian neurogenesis that we define here (i.e. Shared LC1 & 2).

**Figure 2:**
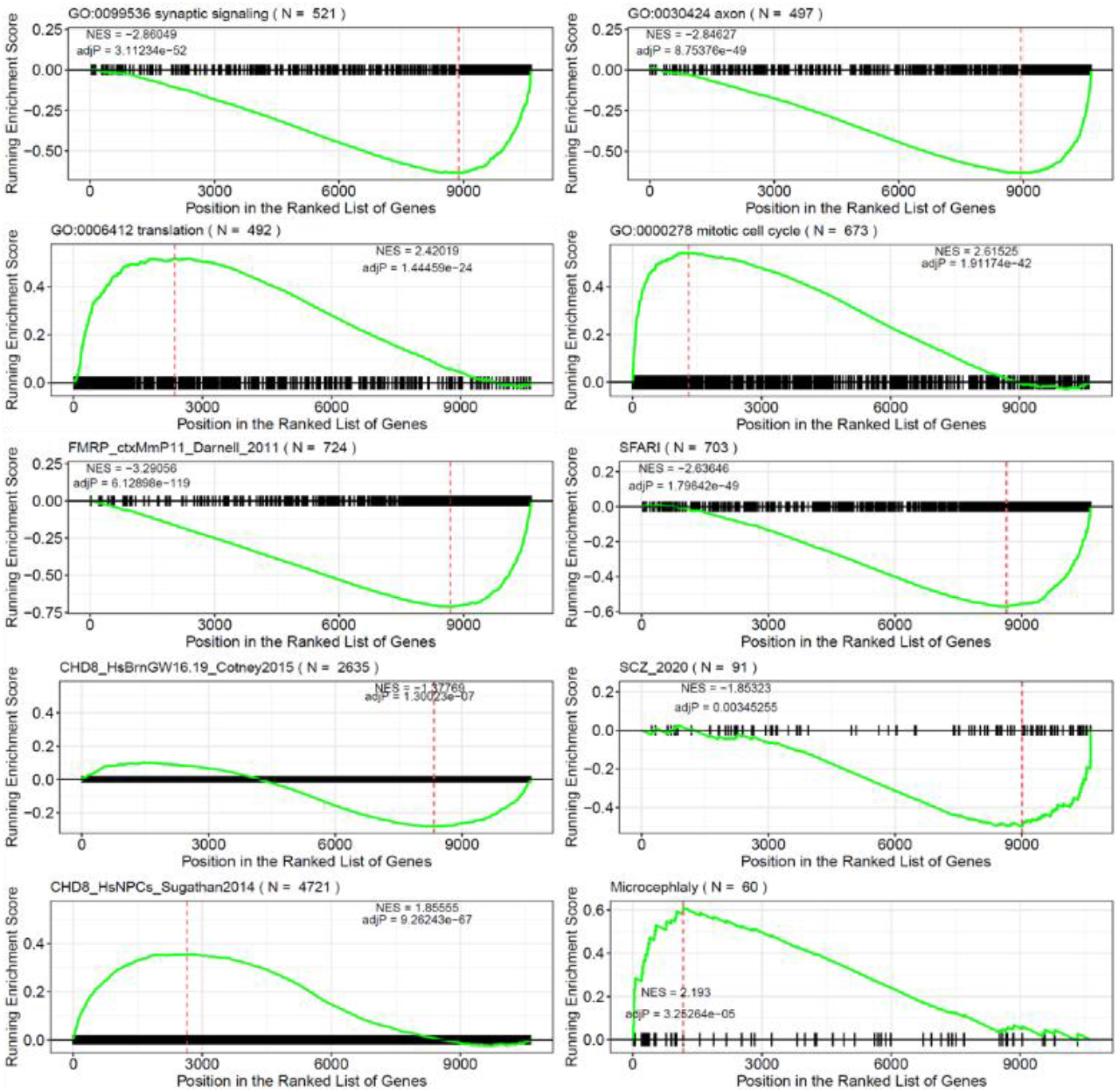
Gene set enrichment analysis (GSEA) on gene loadings associated with LC1 shared across all 4 matrices in Figure 1A. Here, for selected gene groups we show the normalized enrichment scores (NES), p-values, and running enrichment scores visualized across the ranked gene loadings from the shared LC1. As seen in Figure 1A, individual neural stem cells (NSCs) have high embeddings/scores in this LC1, while neurons have low values. The gene loadings reflect this functional cellular distinction across this component of variation: Gene Ontology terms (GO; http://www.gseamsigdb.org/gsea/msigdb/collections.jsp) show the clear enrichment of genes involved in neuronal elements, including the synapse and axon, at the low (neuronal) end of the gene loadings (first row of panels, seen as a right and downward shift in the green running enrichment scores), while genes involved in translation and the cell cycle are enriched at the high (NSC) end (second row of panels, left and up in green). Genes that have been associated with risk for autism spectrum disorder (SFARI; https://gene.sfari.org) and schizophrenia (SCZ; Ripke et al 2020) showed enrichment at the neuronal end of LC1, while genes involved in microcephaly (Jayaraman et al 2018) function more specifically in NSCs. Targets of the autism-associated RNA-binding FMRP defined in the mouse brain (Darnel et al 2011) and targets of autism-associated chromatin regulating CDH8 defined in a mix of cell types in the human brain (Cotney et al 2015) were both enriched at the neuronal end of this trajectory. Interestingly, targets of CHD8 defined in iPSC-derived NSCs grown *in vitro* (Sugathan et al 2014) showed enrichment in the NSC end of this dimension, demonstrating that genes conferring risk for neurodevelopmental and neuropsychiatric disease may do so in multiple cell types at distinct developmental stages. The clusterProfiler (Wu et al 2021) and fgsea (Korotkevich et al 2019) packages in R was used for this GSEA analysis.

Despite this, 2sLCA was able to distinguish pluripotent cells in the *in vitro* data within expression dimensions orthogonal to the shared LC1 (Figure 1B). The ability of this *in vitro* LC to identify pluripotent cells was validated via projection into bulk RNA-seq data from another *in vitro* study that included both pluripotency and differentiation to the neural lineage (Micali et al 2020; Figure 1C).

This demonstrates that 2sLCA was able to correctly dissect a signal known to be common to all the datasets, as well as an element known to be specific to a subset of the matrices being decomposed. Novel in this result is the definition of transcriptome dynamics in cortical neurogenesis that are explicitly 1] common to both human and mouse, and 2] present *in vivo* and recapitulated *in vitro* (see Supplemental Table 1 for gene specific loadings).

## Discussion

With the goal of more fully harnessing the combined discovery power of existing and emerging mutli-omics data across the many fields of biomedicine, we have developed SJD, an R package designed to identify conserved molecular variation across diverse cellular systems. To demonstrate the generality of our approach and its flexibility in harnessing available data, we performed a decomposition of a collection of public gene expression matrices (each processed independently by the original authors), which span *in vivo & in vitro* systems, bulk & scRNA-seq technologies, and mouse & human species. These methods are not proposed as a substitute for tools that define unified dimension reduced spaces and cell type classifications. Instead, it is complementary, defining shared variation that may span spaces and cell clusters defined by these other tools, e.g. transcriptional programs common to multiple distinct cell types. While gene and cell clustering methods define canonical gene function and cell types, our approaches aim to expand exploration of multi-omic data and probe the complex pleiotropy known to underlie genome and cellular function.

## Supplemental Material

### 1-Data normalization in the SJD decomposition functions

Simple data normalization is carried out within each of the decomposition functions in SJD prior to the decomposition itself. With the exception of the jointNMF() method, in all SJD decomposition methods rows/features/genes are set to have mean=0 and columns/samples/cells are set to have mean=0 and standard deviation=1. The row centering is particularly important for defining relative latent spaces in the two-stage methods, twoStageLCA() and twoStageICA(), which perform decomposition on the result of PCA or ICA on the original data (Chen et. al. 2020).

While the column scaling does not ensure a normal distribution of values within each sample, the aggregate distribution of values across columns will be closer to a normal distribution following column scaling. This is a factor that has largely been overlooked in many current methods which apply PCA to non-normally distributed scRNA-seq data, and is the motivation behind the recently developed generalized principal component analysis (GLM-PCA), proposed as a significant improvement over processing non-normal data with conventional PCA (Townes FW, Hicks SC, Aryee MJ, Irizarry RA. Feature selection and dimension reduction for single-cell RNA-Seq based on a multinomial model. Genome Biol. 2019 Dec 23;20(1):295. doi: 10.1186/s13059-019-1861-6. PMID: 31870412.).

In the example we use in this report, we have run SJD on column normalized, log2 transformed, gene level RNA-seq counts. By mean centering on the log scale (the default normalization as described above), we are effectively scaling the raw numbers, i.e. log(x) - mean(log(x)) = log( x/ exp(mean(log(x))), that is, x scaled by the geometric mean. Users can alter the normalization steps carried out on data prior to SJD decompositions by using the rowCenter, rowScale, colCenter, and colScale arguments. While these decomposition functions are designed to be run on a variety of levels of raw or processed data, we recommend an identical or similar normalization process across the matrices used as input in any one SJD decomposition.

As stated above, the one exception to these normalization procedures is in the jointNMF() function, which omits row centering to avoid negative values. Additionally, in all the “joint” analyses (and the concatenation analyses described below), each matrix is divided by the square root of the number of samples in that matrix, with the aim of making the total variation between different data sets comparable (this is not necessary in the two-Stage methods).

Dropping low expression values has always been an important step in the quality control and processing of scRNA-seq data and is generally done by dropping entire rows/genes from an individual matrix if signal is too low. This is even more critical when row normalization is being applied. For example, in the Seurat data integration pipeline (Hao et. al. 2019) where row normalization is also used, only the top 2,000 most variable rows are used by default. This is particularly necessary in the case of Seurat as row variance scaling is also performed. While we do not use row scaling in any of our decompositions and therefore do not recommend dropping such a large a proportion of the data in SJD analyses, we do caution users to drop rows of the data which may contain especially low signal to noise ratio, because even the row centering that SJD does perform can magnify such noise. This comes with the caveat that dropping a row/gene from any of the multiple matrices sent to an SJD joint decomposition will drop that row from the entire analysis. As pointed out in the discussion of pluripotency in the main text (Figure 1B and related text), if a cell type or cellular process is not present in one of the matrices included in a joint decomposition, this can reduce the ability of the decomposition to identify dynamics within that cellular subspace, even if it is present in the other matrices. This should be kept in mind as users assemble related, but diverse matrices for joint decomposition. The “screen_prob” argument in the SJD decomposition functions can be set in order to automatically drop rows/genes lying below a certain proportion of row/gene standard deviations within each matrix (i.e. rows are dropped from each matrix using the within matrix standard deviation and then only common rows kept across all matrices).

### 2 Mathematical details of the “joint” methods as distinct from the two-stage methods

The joint method is focused on the following approximation of the *joint matrix X*:

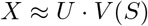

Here, *X* ∈ *R*^*p*×*N*^ is the column stack of multiple input matrices, *U* ∈ *R*^*p*×*r*^ is the components matrix (with *r* components), *V* ∈ *R*^*r*×*N*^ is the score matrix, and *S* ∈ *R*^*r*×*N*^ is the structural matrix, which encodes the structure among those multiple data sets.

The symbol “≈” indicates approximation, which is crucial in determining the final learned components and structure of the model. We propose three different ways to achieve this approximation: **principal component analysis (PCA), independent component analysis (ICA)**, and **nonnegative matrix factorization (NMF)**.

#### Principal Component Analysis (PCA)

For the PCA method, we optimize the following objective function:

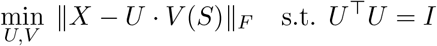

To estimate this objective, we use a two-stage method:

1. Randomly initialize *U* and *V* with the corresponding fixed dimensions;
2. Fix *U*, and use least squares to compute *V* ;
3. Fix *V*, and use the Procrustes algorithm to compute *U* under the orthogonal constraint;
4. Iterate between steps 2 and 3 until convergence.

#### Independent Component Analysis (ICA)

For the ICA method, we optimize:

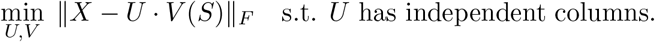

We compute this based on the output of the PCA method. We apply the FastICA algorithm on the matrix

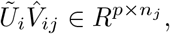

which is the extracted data from the *j*-th dataset based on the *i*-th component.

#### Nonnegative Matrix Factorization (NMF)

For the NMF method, we optimize:

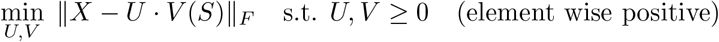

To estimate this objective, we use a two-stage method:

1. Randomly initialize *U* and *V* to be nonnegative;
2. Fix *U*, optimize *V* with the nonnegativity constraint;
3. Fix *V*, optimize *U* with the nonnegativity constraint;
4. Iterate between steps 2 and 3 until convergence.

#### ProjectiLCA: ICA-based projection

The projectiLCA() function projects a new dataset *D* into an existing ICA-based decomposition from twoStageiLCA(). It takes as input a new data matrix D, gene score matrices from the original iLCA decomposition, and ICA results (pre-whitening matrix *K* and unmixing matrix *W* ) from the original iLCA decomposition.

In projection:

The new data *D* is first multiplied by the LCA gene scores *G* to obtain the projected LCA sample score matrix *S*_*a*_:

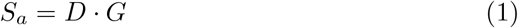

Then the LCA sample score matrix *S*_*a*_ is multiplied by the pre-whiting matrix *K* and the unmixing matrix *W* :

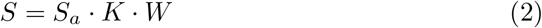

And *S* is the resulting sample score from the iLCA projection.

This two-step projection extracts the independent components of the new data while maintaining consistency with the original ICA decomposition.

### 3– Further notes on differences between two-stage and the “joint” methods

The two-stage methods twoStageLCA() and twoStageiLCA(), as well as the “joint” decomposition analyses jointPCA(), jointICA(), and jointNMF() in SJD all consider the hierarchical structure between datasets that is defined by the user via the matrix groupings argument. Because the “joint” methods estimate all requested components simultaneously, it is possible that when components have significant differences in variance across groupings, this method will produce sub-optimal separation of effects. The two-stage methods estimate components in the matrix grouping order specified by the user and sequentially search spaces orthogonal to previous groupings’ components. This allows twoStageLCA() and twoStageiLCA() to be more computationally efficient and to achieve greater separation of components across matrix groupings than the “joint” methods, especially when differences in variance across groupings is large. It should be noted that while these methods are based on the math of conventional PCA and ICA, the resulting LCs are not variance ranked as are the component resulting from the conventional, individual matrix analyses.

### 4-Additional functionality of the SJD decomposition functions

While we have requested a small number of components in the example application illustrated in main Figure 1, the decomposition tools in the SJD package are also intended define higher dimensional shared latent spaces across datasets. We have successfully used SJD to identify 10’s of biologically relevant components shared across matrices. When requesting higher numbers of components that span groups of matrices, users should keep in mind the fundamentally different decompositions offered by PCA (used in twoStageLCA() and jointPCA()), ICA (used in twoStageICA() and jointICA()), and NMF (used in jointNMF()).

Additionally, the SJD decomposition functions include two capabilities not discussed in the main text of the manuscript:

1. Differential weighting of individual matrices in the decompositions, using the “weighting” argument. This can be used in many ways, but is particularly useful if there are many studies of one kind and a different number of studies of a second kind, but the user wants the total weighting of the two kinds of study to be equal. For example, if the user desires a decomposition that equally weighs a collection of matrices that includes both *in vivo* experiments and *in vitro* experiments, but there are differing numbers of the 2 kinds of experiment. In this case, the user could give equal weights to the *in vivo* studies that sum to 1 (i.e. weight=1/N *in vivo* studies) and similarly weights to the *in vitro* studies that sum to 1 (i.e. weight=1/N *in vitro* studies), giving *in vivo* and *in vitro* data equal weighting regardless of the number of *in vivo* and *in vitro* studies.
2. In addition to the data matrices that the user wants included in the decomposition, the user can also include data matrices in SJD function calls that are not intended to affect the decomposition, but in which the user wants to see the components of variation. Using the sjdWrapProjection() function the rows of the projection only matrices can be aligned (as is done with the sjdWrap() function for matrices being included in the joint decomposition). The “proj_dataset” argument in the SJD decomposition functions takes the list of matrices to be used for projection only and the “proj_group” argument is used to indicate components (i.e. which matrix groupings) onto which each dataset is requested be projected, i.e. a list of length equal to the number of matrices in the “proj_dataset” argument, where each of these list elements contains a logical of length equal to the length of the “group” argument.

### 5-BEMA rank determination in the “twoStageLCA” function

In addition to the “twoStageLCA” function which is fixed rank and requires the user to input the fixed dimensions for each component, the SJD package also offers the function “twoStageLCA.rank” which executes rank selection as part of the 2sLCA method where the dimensionality of each component space is determined automatically. Like the “twoStageLCA” function, the user is required to enter both the datasets and grouping lists, however, the difference lies in that rather than entering the number of components for each group, the user needs to enter the “threshold” for each component (e.g. if the component is shared by 4 datasets and the next component is shared by only 2 datasets, then a recommended threshold should be 3, as it lies between 2 and 4, and the algorithm will be able to detect the gap between 2 and 4). The “twoStageLCA.rank” function returns output with the same structure as the output of the “twoStageLCA” function.

The method of two-stage linked component analysis (2sLCA) is built on principal component analysis (PCA). First, for each dataset, PCA is conducted and the principal spaces associated with top singular values are extracted. In this step, the dimensions of the principal spaces are determined via the BEMA method (KE, Z. T., MA, Y. AND LIN, X. 2021. Estimation of the number of spiked eigenvalues in a covariance matrix by bulk eigenvalue matching analysis. *Journal of the American Statistical Association* 1–19), which detects signals from noise by fitting a distribution to the observed bulk eigenvalues arising from noise. Second, all the principal spaces are combined to extract the common subspace, the partially shared subspaces and the individual subspaces, also using PCA. The subspaces are assumed orthogonal to each other to ensure their identifiability. Moreover, the dimensions of the subspaces can be determined by the natural gaps of eigenvalues associated with the subspaces. This procedure is theoretically valid if the subspaces shared by more datasets are first extracted.

### 6-Additional decomposition functions for testing and comparison

Additional functions to conduct dimension reduction on individual matrices, i.e. sepPCA(), sepICA() and sepNMF() and concatenated families of matrices, i.e. concatPCA(), concatICA() and concatNMF() have also been included in SJD for comparative analyses. Although common in the literature, we do not recommend dimension reduction on concatenated matrices using PCA or ICA approaches unless the datasets used are known to be co-registered from an experimental point of view, i.e. identical sample collection, RNA extraction, library preparation, sequencing protocol, and data processing within a single laboratory. Even in this case, using the concatenation methods may result in greater overlap between partially shared components and an unintentional focus on technical artifacts that exist between the original matrices. For this same reason, if NMF approaches are applied to concatenated matrices (as is the case when using either concatNMF() or when using jointNMF() while requesting only dimensions shared by all matrices, see PMID: 38464021 for examples) care should be taken to explore differences in sample/cell embeddings/scores only WITHIN matrices/experiments.

The SJD package contains skreeCompare() and sjd2tSNEumap() functions that enable users to compare SJD’s joint decomposition results with conventional individual matrix decomposition. In addition, the SJDScorePlotter(), assemble.byDataset(), and assemble.byComponent() functions allow the ordered visualization of the many related dimensions that SJD analyses can produce across large families of matrices. Code implementing these functions can be found at: https://github.com/carlocolantuoni/NeocortexDevelopment_Sonthalia2024/blob/main/code/SJDcode_Figure1n2n3.R.

### 7-Other methods that relate to our work on SJD

*In general these approaches have more specific data type utility and analytical goals (e*.*g. “integration” methods that seek to map multiple datasets of specific modalities onto a unified dimension reduced space) than SJD, which we have designed to be a broadly applicable joint dimension reduction tool that can be applied to any matrices with shared measurements/features (e*.*g. genes, genomic coordinates, spatial locations, etc*.*) across samples/observations, and subsequently combined with other exploratory approaches*.

